# Pathophysiological Implications of Nucleotide Self-Assembly: Adenine-Derived Nucleotides Aggregation in Disease Mechanisms

**DOI:** 10.1101/2024.10.28.620770

**Authors:** Raj Dave, Ankur Singh, Kshipra Pandey, Ritu Patel, Nidhi Gour, Dhiraj Bhatia

## Abstract

Adenine nucleotides, including adenosine monophosphate, adenosine diphosphate, and adenosine triphosphate, play pivotal roles in cellular bioenergetics, nucleic acid metabolism, and signal transduction. However, their propensity to undergo self-assembly and form supramolecular aggregates under certain conditions is not well-characterized. In this study, we examined the self-assembly, aggregation, and cytotoxicity of AMP, ADP, and ATP in both fresh and aged conditions. Utilizing advanced microscopy techniques, Thioflavin T (ThT) fluorescence assays, and cross-seeding experiments, we identified oligomer formation in freshly prepared nucleotide solutions, which progressed to larger, more stable aggregates over time. The cytotoxic potential of these nucleotide aggregates was assessed using in vitro models, including human retinal pigment epithelial (RPE-1) and colorectal carcinoma (HCT-116) cell lines. Our findings demonstrate that nucleotide aggregation induces significant cytotoxic effects, particularly in aged conditions. Further investigations into bacterial toxicity models revealed similar deleterious impacts, indicating a broad-spectrum biological stress response to nucleotide aggregates. These results suggest that nucleotide self-assembly and aggregation may contribute to cellular dysfunction, offering new insights into their non-canonical roles in disease pathophysiology, potentially analogous to protein misfolding disorders.

## Introduction

Nucleotides are fundamental molecules involved in the essential functions of living organisms. ATP, for instance, is the primary energy currency of the cell, driving numerous enzymatic reactions, including those involved in metabolism, muscle contraction, and cellular signaling [1]. AMP and ADP, on the other hand, are involved in cellular energy homeostasis and act as intermediates in the regeneration of ATP via the enzymatic action of ATP synthase. Additionally, nucleotides serve as building blocks for nucleic acids, playing a crucial role in DNA and RNA synthesis, which is vital for cell replication and repair [2]. Furthermore, nucleotides are involved in signaling pathways such as purinergic signaling, where extracellular ATP acts as a signaling molecule, mediating processes like inflammation, vasodilation, and neurotransmission [3]. Imbalances in nucleotide metabolism or their extracellular accumulation can lead to pathological conditions, including energy metabolism disorders and inflammatory responses [4]. Understanding the behavior of nucleotides, particularly in their aggregated form, could provide novel insights into their involvement in disease states. Beyond their canonical roles, nucleotides can also exhibit dynamic structural behaviors, particularly under certain conditions, where they have the ability to self-assemble into higher-order structures. This self-assembly process can lead to the formation of oligomers and, over time, larger aggregates, which may impact cellular functions [5]. Recent studies have highlighted the potential for nucleotide aggregation to result in cytotoxicity, a phenomenon commonly observed in the context of protein aggregation, such as in neurodegenerative diseases [6]. However, the self-assembly and aggregation of nucleotides and their toxicological implications remain an overlooked area of study. Understanding how these molecules transit from monomers to oligomers and ultimately to aggregated forms under both fresh and aged conditions could provide insight into previously unexplored mechanisms of cellular dysfunction and toxicity. In this study, we explore the self-assembly and aggregation of AMP, ADP, and ATP under fresh and aged conditions using a range of biophysical and biological assays. By employing microscopy techniques, Thioflavin T (ThT) binding assays, and cross-seeding experiments, we have visualized the oligomeric and aggregated forms of these nucleotides. Moreover, we extended our investigation to in vitro systems, using RPE-1 and HCT-116 cell lines, to assess the cytotoxicity of these aggregates. Additionally, the interactions between nucleotide aggregates and bacterial cells were explored, revealing significant toxicity. The formation of oligomers in fresh nucleotide solutions and the development of more stable aggregates with aging point to a complex mechanism of nucleotide self-assembly that could have broader biological implications. This work aims to build on the hypothesis that nucleotide aggregation, similar to protein aggregation, may contribute to cellular stress and toxicity, thus providing new insights into nucleotide behaviour under physiological and pathological conditions.

## Result and Discussion

The self-assembly behavior of AMP demonstrates its amyloidogenic potential, as observed through multiple techniques including optical microscopy, SEM, and AFM. Initially, AMP forms oligomeric species, but with aging, these aggregates mature into a fibrillar network, indicative of amyloid fibril formation [7]. The increase in ThT fluorescence over time further supports the progression of AMP into β-sheet-rich structures, characteristic of amyloid fibrils [8]. Optical microscopy images (Figure 2A) revealed the progression of AMP (1 mM) self-assembly over time. On Day 1 (Figure 2A, i), oligomeric structures were predominant, indicating early-stage aggregation [9]. By Day 5 (Figure 2A, ii), the presence of fiber-like assemblies began to emerge, and by Day 10 (Figure 2A, iii), a well-defined network of fibrils was observed, suggesting the amyloidogenic nature of AMP. To further confirm the amyloidogenic properties of AMP, a ThT binding assay was performed [10] (Figure 2B). On Day 1 (Figure 2B, i), minimal fluorescence was observed, indicating a low level of β-sheet formation. By Day 5 (Figure 2B, ii), ThT fluorescence increased significantly, correlating with the formation of structured fibrils. This trend was even more pronounced on Day 10 (Figure 2B, iii), confirming the progression of AMP into amyloid-like fibers.

**Figure 1.**
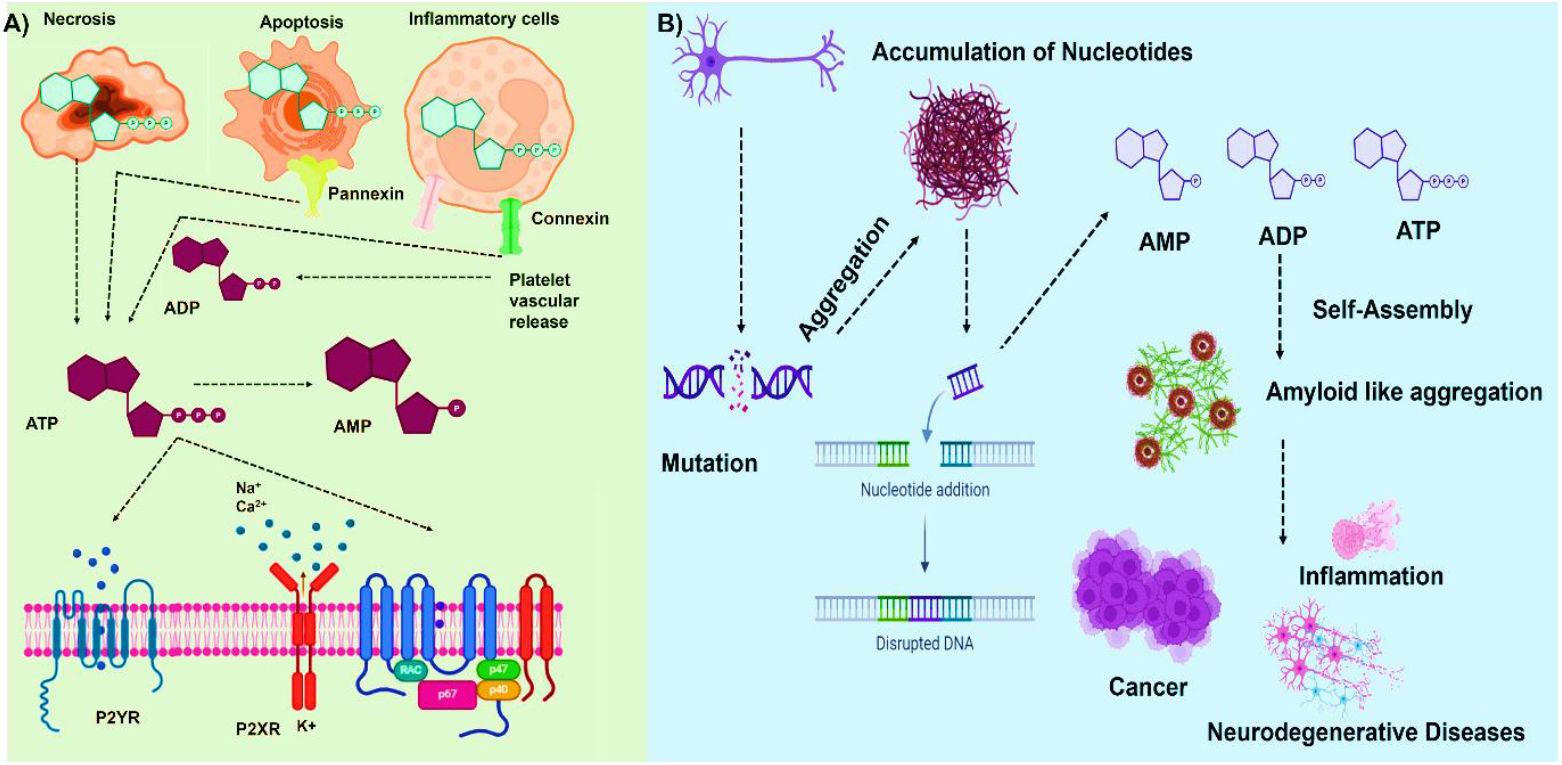
Schematic representation of accumulation of AMP, ADP and ATP caused by gene mutation and consequence, which lead to rare IEMs, cancer, neurodegenerative diseases etc.

**Figure 2.**
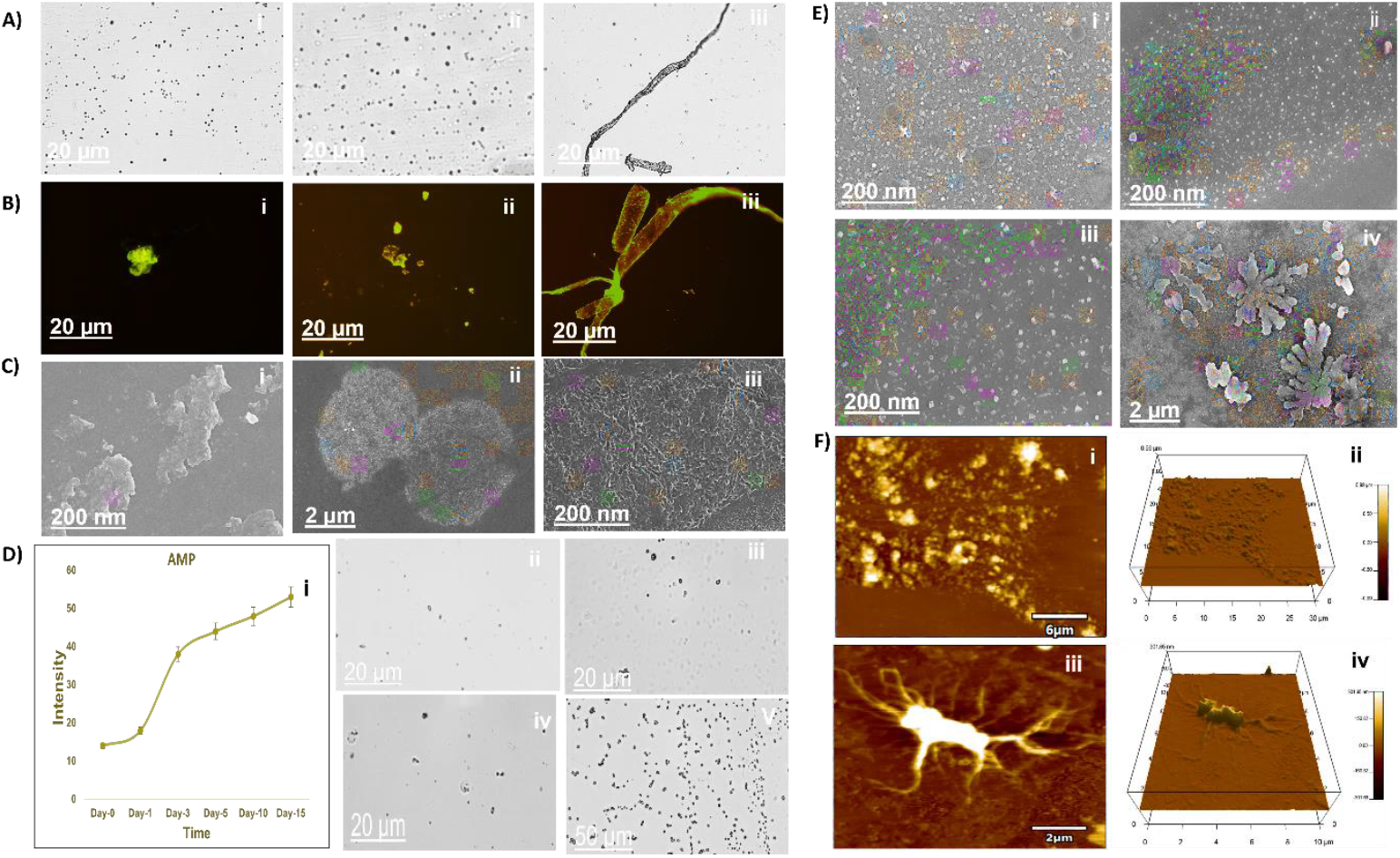
Self-assembly of AMP (1mM) (A) Optical microscopy of AMP at different time intervals; (i, ii and iii) Day-1, Day-5 and Day-10; (B) ThT binding assay of AMP; (i, ii and iii) Day-1, Day-5 and Day-10;(C) Cross seeding experiments; (i, ii and iii) AMP alone, AMP co-incubated with urea and AMP co-incubated with tannic acid; (D) (i) AMP at solution state, (ii-v)AMP at pH 3,6,8 and 10; (E) SEM microscopy at different time intervals; (i, ii) Fresh conditions (iii, iv) Aged conditions and (F) AFM microscopy at different time intervals; (i, ii) 2D and 3D AFM micrograph of AMP at fresh conditions (iii, iv) 2D and 3D AFM micrograph of AMP at aged conditions.

The ability of urea and tannic acid to disassemble these aggregates highlights the importance of hydrogen bonding in maintaining the fibrillar structure of AMP. Urea, as a chaotropic agent, disrupts hydrogen bonds, leading to the breakdown of the fiber-like assemblies. Similarly, tannic acid disrupts the aggregation, reinforcing its potential role as an anti-amyloidogenic agent [11]. These findings open potential therapeutic avenues for the prevention of amyloid-related diseases. Urea, a known hydrogen bond disruptor, effectively disassembled the AMP aggregates (Figure 2C, ii, SI 1). Similarly, tannic acid, which interferes with protein aggregation, also showed a strong capacity to break the fiber-like network (Figure 2C, iii, SI 2). These results confirm that both compounds effectively disrupt the hydrogen-bonded structures of AMP fibrils. AMP was tested under various pH conditions to investigate its structural stability in different buffer systems (Figure 2D). At pH 3, 6, 8, and 10 (Figure 2D, ii-v), AMP assemblies exhibited salt crystal formation, suggesting a significant influence of pH on the self-assembly process. The pH variation did not show any significant fibril formation but rather induced salt crystallization in most conditions. Additionally, the pH-dependent behavior of AMP suggests that the self-assembly process can be significantly influenced by environmental conditions. The observation of salt crystal formation at various pH levels indicate that AMP’s aggregation pathway is sensitive to ionic strength and pH, further supporting the role of buffer conditions in modulating protein assembly [12]. Scanning electron microscopy (SEM) was used to assess the morphological changes in AMP over time [13] (Figure 2E). Fresh samples (Figure 2E, i and ii) primarily showed monomeric or small oligomeric units. In contrast, aged samples (Figure 2E, iii and iv) exhibited the formation of a dense network of fibers, consistent with the optical microscopy observations. Atomic force microscopy (AFM) provided high-resolution topographical images of AMP aggregates. In fresh conditions (Figure 2F, i and ii), AFM revealed small, round monomers and oligomers with minimal network formation. After aging (Figure 2F, iii and iv), the AFM micrographs displayed a well-defined, interconnected fibrillar network, consistent with the observations from both SEM and optical microscopy. The consistent findings across optical microscopy, SEM, and AFM provide robust evidence of AMP’s self-assembly dynamics and its transition from monomers to amyloid-like fibrils [14]. Understanding these pathways is crucial for designing interventions that target or modulate amyloid formation in pathological conditions.

The self-assembly behavior of ADP (1 mM) was explored over time using multiple techniques, starting with optical microscopy (Figure 3A). On Day 1 (Figure 3A, i), ADP primarily existed as dispersed monomeric or oligomeric units, indicating an early stage in the aggregation process. By Day 5 (Figure 3A, ii), fiber-like structures began to emerge, suggesting intermediate aggregation. On Day 10 (Figure 3A, iii), the network of fibrillar structures became more prominent, indicating a mature assembly state. These temporal changes illustrate the gradual transition of ADP from an initial monomeric state to a more organized, amyloid-like fibrillar structure. The amyloidogenic properties of ADP were further assessed using the Thioflavin T (ThT) binding assay (Figure 3B). On Day 1 (Figure 3B, i), minimal fluorescence was detected, suggesting limited β-sheet formation at the early stage. However, by Day 5 (Figure 3B, ii), ThT fluorescence increased significantly, corresponding to the formation of ordered aggregates. This fluorescence intensity peaked by Day 10 (Figure 3B, iii), confirming the development of amyloid-like fibrils in the ADP solution. In cross-seeding experiments, the role of urea and tannic acid in disrupting ADP assemblies was investigated (Figure 3C). ADP incubated alone (Figure 3C, i) formed well-defined fibers, whereas co-incubation with urea, a hydrogen bond disruptor, led to a significant reduction in fibril formation (Figure 3C, ii, SI 1). Similarly, tannic acid, known for its anti-aggregation properties, was effective in breaking down the ADP fibers (Figure 3C, iii SI 2). These observations highlight the critical role of hydrogen bonding in stabilizing ADP aggregates and the ability of specific agents to interfere with this process. The effect of pH on ADP self-assembly was studied using different buffer systems (Figure 3D). ADP in solution state (Figure 3D, i) displayed no significant aggregation. However, at varying pH levels (3, 6, 8, and 10) (Figure 3D, ii-v), ADP showed the formation of salt crystals, particularly at extreme pH values, indicating that environmental conditions, such as pH, play a significant role in influencing ADP aggregation pathways. Despite these changes, no pronounced fibril formation was detected across the pH conditions tested, emphasizing that ADP’s self-assembly is highly sensitive to pH and ionic strength. Scanning electron microscopy (SEM) provided a detailed view of the morphological changes in ADP over time (Figure 3E). Freshly prepared ADP samples (Figure 3E, i and ii) primarily exhibited small, monomeric or oligomeric structures, with minimal organization. In contrast, aged samples (Figure 3E, iii and iv) displayed a robust network of fibrils, consistent with the aggregation observed in optical microscopy and ThT binding assays. Atomic force microscopy (AFM) offered additional insights into the topographical features of ADP aggregates (Figure 3F). Fresh ADP samples (Figure 3F, i and ii) revealed small, scattered aggregates, while aged samples (Figure 3F, iii and iv) showed extensive fibrillar networks with significant height variation, indicating a progression from monomeric units to structured fibrils over time. The 3D AFM images confirmed the dense, interconnected nature of these fibrils in aged samples. Taken together, these results reveal a clear progression of ADP from monomers to amyloid-like fibrils and demonstrate the impact of environmental factors and specific inhibitors on this process. This study provides a foundation for future research into the mechanisms of ADP aggregation and its potential implications for amyloid-related diseases.

**Figure 3.**
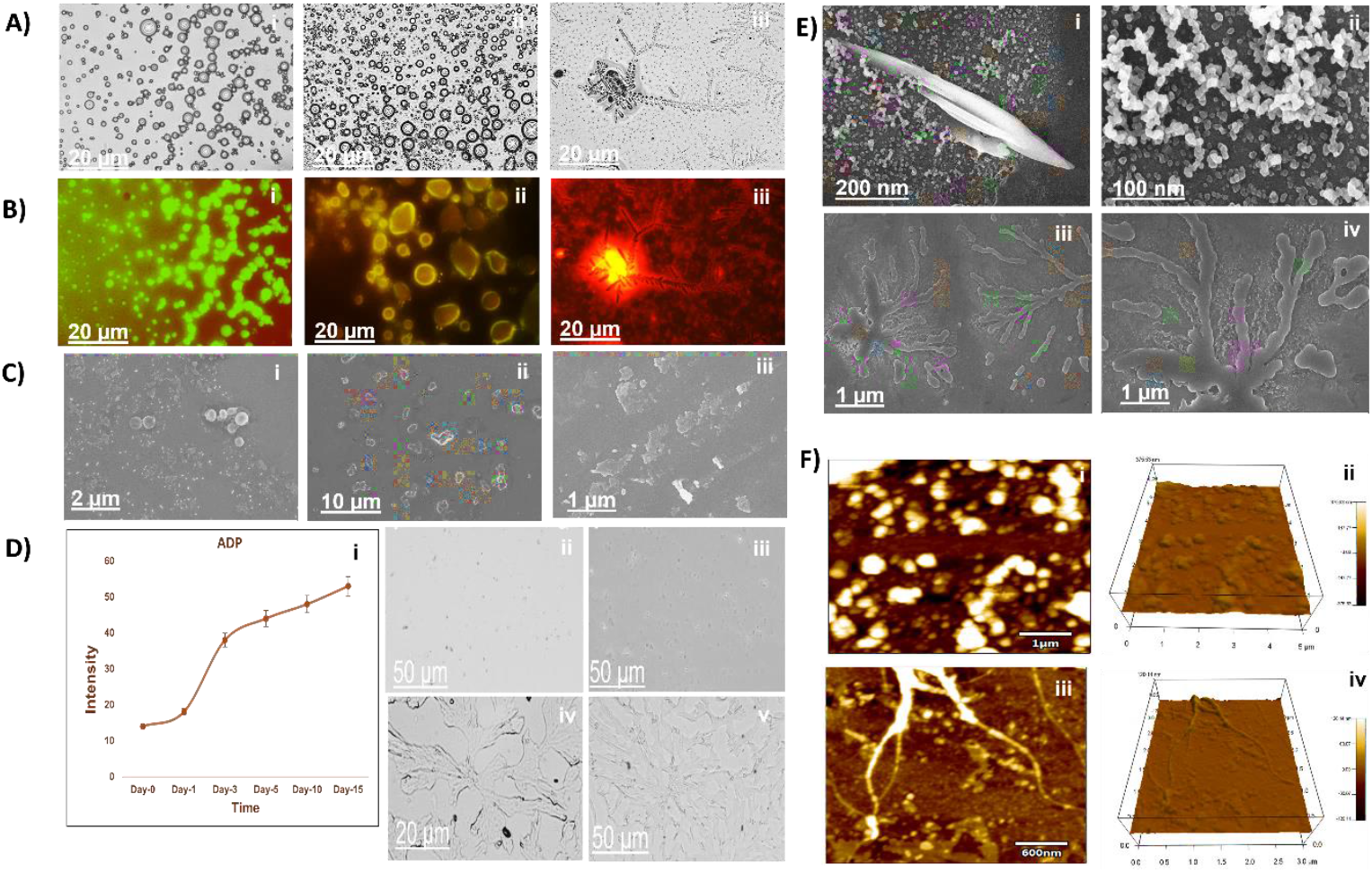
Self-assembly of ADP (1mM) (A) Optical microscopy of ADP at different time intervals; (i, ii and iii) Day-1, Day-5 and Day-10; (B) ThT binding assay of ADP; (i, ii and iii) Day-1, Day-5 and Day-10;(C) Cross seeding experiments; (i, ii and iii) ADP alone, ADP co-incubated with urea and ADP co-incubated with tannic acid; (D) (i) ADP at solution state, (ii-v)ADP at pH 3,6,8 and 10; (E) SEM microscopy at different time intervals; (i, ii) Fresh conditions (iii, iv) Aged conditions and (F) AFM microscopy at different time intervals; (i, ii) 2D and 3D AFM micrograph of ADP at fresh conditions (iii, iv) 2D and 3D AFM micrograph of ADP at aged conditions.

The self-assembly of ATP (1 mM) was systematically evaluated over time using multiple analytical techniques. Initial optical microscopy (Figure 4A) revealed distinct morphological transitions. On Day 1 (Figure 4A, i), ATP predominantly existed as dispersed monomeric or small oligomeric units, characteristic of early nucleation events. By Day 5 (Figure 4A, ii), an increase in organized fibrillar structures was observed, indicating the formation of protofibrils. By Day 10 (Figure 4A, iii), these protofibrils had matured into an extensive fibrillar network, suggesting a transition toward a stable amyloid-like assembly over time. The amyloidogenic propensity of ATP was further corroborated by the Thioflavin T (ThT) fluorescence assay (Figure 4B). On Day 1 (Figure 4B, i), minimal ThT fluorescence was detected, indicating a low degree of β-sheet content. However, by Day 5 (Figure 4B, ii), a notable increase in fluorescence intensity was observed, suggesting the formation of ordered β-sheet-rich aggregates. By Day 10 (Figure 4B, iii), the ThT fluorescence plateaued at a high intensity, confirming the accumulation of stable amyloid fibrils. In cross-seeding experiments, the effect of co-incubation with urea and tannic acid on ATP self-assembly was investigated (Figure 4C). ATP incubated alone (Figure 4C, i) formed a well-structured fibrillar network. However, when co-incubated with urea, a chaotropic agent that disrupts hydrogen bonds, significant disassembly of the fibrils was observed (Figure 4C, ii, SI 1), suggesting that hydrogen bonding plays a crucial role in the stabilization of ATP fibrils. Similarly, tannic acid, known for its anti-amyloidogenic properties, also demonstrated effective disruption of the ATP fibrillar network (Figure 4C, iii, SI 2), confirming its potential in inhibiting ATP aggregation. The influence of pH on ATP aggregation was assessed across different buffer systems (pH 3, 6, 8, and 10) (Figure 4D). In the solution state (Figure 4D, i), ATP displayed no significant aggregation. At pH 3 and 6 (Figure 4D, ii and iii), the formation of salt crystals was observed, likely due to increased ionic interactions in these acidic environments. At pH 8 and 10 (Figure 4D, iv and v), ATP exhibited reduced salt crystal formation but showed minimal fibril organization, suggesting that extreme pH conditions hinder the self-assembly process, potentially by disrupting critical intermolecular interactions. Scanning electron microscopy (SEM) provided detailed insights into the morphological transitions of ATP over time (Figure 4E). Fresh samples (Figure 4E, i and ii) displayed dispersed monomeric and oligomeric species with no defined structure. In contrast, aged samples (Figure 4E, iii and iv) exhibited extensive fibrillar networks, consistent with the progression from initial nucleation to mature amyloid-like fibrils observed in optical microscopy and ThT binding assays. Atomic force microscopy (AFM) was employed to further characterize the topographical features of ATP aggregates (Figure 4F). Fresh ATP samples (Figure 4F, i and ii) displayed amorphous, small aggregates with minimal height variation, indicative of early-stage assembly. However, aged ATP samples (Figure 4F, iii and iv) revealed a dense network of mature fibrils, with significant structural complexity and height, confirming the formation of highly organized fibrillar assemblies over time. Overall, this study provides a comprehensive characterization of ATP self-assembly, revealing its amyloidogenic nature and the key factors that influence its aggregation. These findings contribute to our understanding of nucleotide-driven aggregation processes, with potential implications for both fundamental science and therapeutic interventions aimed at modulating amyloid formation.

**Figure 4.**
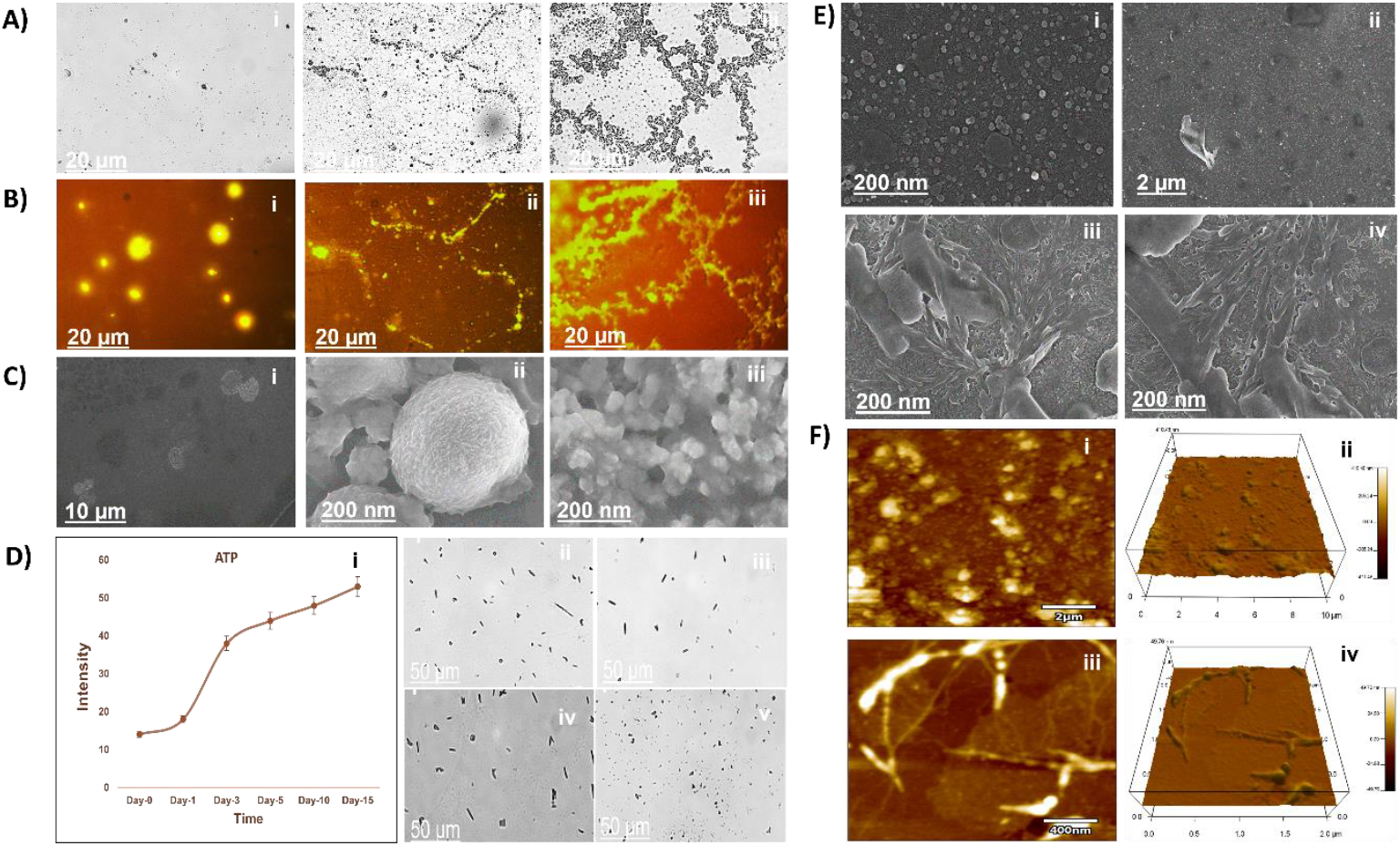
Self-assembly of ATP (1mM) (A) Optical microscopy of ATP at different time intervals; (i, ii and iii) Day-1, Day-5 and Day-10; (B) ThT binding assay of ATP; (i, ii and iii) Day-1, Day-5 and Day-10;(C) Cross seeding experiments; (i, ii and iii) ATP alone, ATP co-incubated with urea and ATP co-incubated with tannic acid; (D) (i) ATP at solution state, (ii-v)ATP at pH 3,6,8 and 10; (E) SEM microscopy at different time intervals; (i, ii) Fresh conditions (iii, iv) Aged conditions and (F) AFM microscopy at different time intervals; (i, ii) 2D and 3D AFM micrograph of ATP at fresh conditions (iii, iv) 2D and 3D AFM micrograph of ATP at aged conditions.

We further investigated the influence of these nucleotide aggregates on cell viability since many amyloids are lethal due to their propensity to damage cell membranes [15]. Thus, using RPE-1 and HCT-116 cells co-incubated with fresh and aged samples of AMP, ADP, and ATP assemblies, we performed the MTT experiment to verify our hypothesis (Figure 5). The MTT assay is a rapid colorimetric test that uses dehydrogenases in active mitochondria to cleave the tetrazolium ring of MTT (3-(4,5-dimethylthazolk-2-yl)-2,5-diphenyl tetrazolium bromide) in order to determine the number of live cells in alive cells [16].

**Figure 5.**
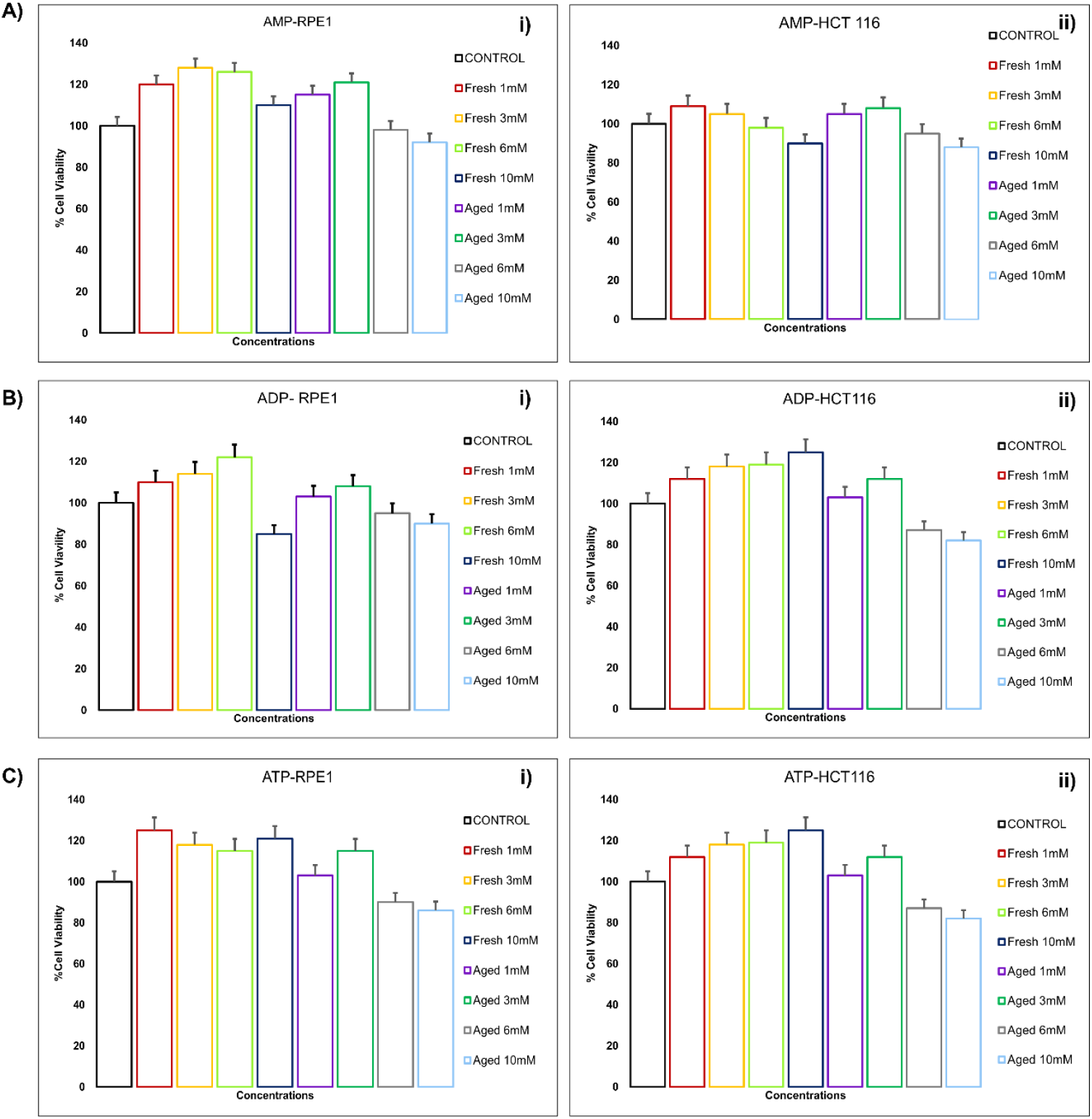
Graphical representation of MTT assay in fresh and in aged conditions when co-incubated in RPE-1 and HCT 116 cell lines with varying concentrations of AMP, ADP and ATP.

The results of the *in vitro* assays for AMP, ADP, and ATP with RPE-1 and HCT116 cells under fresh and aged conditions reveal distinct patterns of cell viability and cytotoxicity. In the case of AMP, when incubated with RPE-1 cells, a clear concentration-dependent effect on cell viability was observed in fresh conditions. At lower concentrations (1 mM and 3 mM), RPE-1 cells maintained high viability, showing no cytotoxic effects. Interestingly, aged AMP, characterized by aggregation, exhibited more pronounced cytotoxic effects at all concentrations, suggesting that the aggregated form of AMP enhances cellular stress or interferes more severely with cellular metabolism. This likely leads to higher oxidative stress or membrane disruption, reducing cell viability more substantially in RPE-1 cells (Figure 5 A, i). In contrast, HCT116 cells exhibited greater resistance to AMP compared to RPE-1 cells. Under fresh conditions, cell viability remained high across all concentrations, with minimal reduction. Even under aged conditions, HCT116 cells demonstrated only a slight reduction in viability at higher AMP concentrations (6 mM and 10 mM), indicating that the cancerous HCT116 cells possess greater resilience to both fresh and aggregated AMP (Figure 5 A, ii). This resistance may be attributed to the altered metabolism and enhanced survival mechanisms in cancer cells, such as upregulated glycolysis or autophagic pathways, which help them withstand cytotoxic stress more effectively than non-cancerous cells [17].

For ADP, the trends observed were similar to AMP. In RPE-1 cells (Figure 5B, i), fresh ADP exhibited the absence of cytotoxic effect, with cell viability increasing as ADP concentration increased. However, under aged conditions, ADP aggregates had a much stronger impact, leading to a significant reduction in viability, particularly at higher concentrations (6 mM and 10 mM). These findings suggest that ADP aggregation may induce further cellular damage, potentially by disrupting energy metabolism or causing oxidative stress. On the other hand, HCT116 cells (Figure 5 B, ii) again demonstrated higher tolerance at fresh conditions whereas the aged ADP demonstrated a cytotoxic effect. This suggests that HCT116 cells also have similar impacts on the cellular pathways assigned to manage the toxic effects of ADP as in RPE-1 cells, possibly due to their compromised metabolic flexibility.

ATP exhibited the most significant cytotoxic effect on RPE-1 cells, particularly under aged conditions. In fresh conditions (Figure 5C, i), ATP caused a notable increase in viability, indicating that excess ATP enhances cellular energy homeostasis. The aged ATP aggregates were more cytotoxic, leading to a sharp decline in viability at lower concentrations (6 mM and 10 mM). This suggests that aged ATP aggregates, like AMP and ADP, may disrupt cell membranes or induce apoptosis through dysregulation of ATP-sensitive pathways. In HCT116 cells (Figure 5C, ii), the cytotoxic effects of ATP were greatly demonstrated when considering the aged condition. Under fresh conditions, ATP caused a significant increase in viability. Aged ATP showed more toxic effects compared to aged ADP and AMP conditions. Aged condition of ATP significantly reduced cell viability, reflecting the enhanced decrease in metabolic adaptability of HCT116 cells, which can mitigate the toxic effects of ATP and its aggregates.

The results demonstrate that RPE-1 cells, a non-cancerous cell line, are more sensitive to the cytotoxic effects of AMP, ADP, and ATP, particularly under aged conditions, where nucleotide aggregation exacerbates the reduction in viability. This could be due to the increased cellular stress induced by nucleotide aggregates, which potentially disrupt cellular membranes or interfere with mitochondrial function. Whereas, HCT116 cells, a cancerous cell line, showed relatively similar results in both fresh and aged forms of AMP, ADP, and ATP. These findings are significant for understanding the effects of nucleotide metabolism in cancerous versus non-cancerous cells and may provide insights into potential therapeutic targets that exploit the metabolic vulnerabilities of normal and cancer cells.

The bacterial study investigated the antimicrobial activity of three nucleotide analogs ATP, ADP, and AMP at varying concentrations (1mM, 3mM, 6mM, and 10mM) against the selected two Gram-positive bacteria (*Bacillus subtilis* and *Staphylococcus aureus*) and two Gram-negative bacteria (*Pseudomonas aeruginosa* and *Vibrio cholerae*) (Figure 6). The data showed distinct bacterial growth responses depending on the concentration and the condition of the test samples, i.e., the freshly prepared condition and the aged condition. Considering the fresh conditions, a concentration-dependent enhancement in bacterial growth was observed. Higher concentrations of the test compounds (6 mM and 10 mM) promoted bacterial proliferation, suggesting that the fresh nucleotide analogs may serve as metabolic substrates, facilitating growth. This growth-promoting effect could be due to the role of nucleotides in cellular energy metabolism, signaling pathways, or nucleotide biosynthesis, which may enhance the bacteria’s overall viability and growth rate at higher doses [18]. In contrast, under the aged conditions, the test samples exhibited a toxic effect on bacterial cultures, resulting in a marked inhibition of growth as the concentration increased. This suggests that over time, the test samples may undergo chemical degradation or structural alterations, generating by-products with cytotoxic properties [19]. These degradation products likely disrupt cellular processes or induce oxidative stress, leading to a reduction in bacterial viability. The differential impact of aged versus fresh samples highlights the importance of stability and chemical integrity when evaluating the biological activity of nucleotide analogs [20]. These findings indicate that ATP, ADP, and AMP exhibit dual roles in bacterial growth modulation: promoting growth under fresh conditions and exerting toxic effects under aged conditions. This suggests that the biological activity of these nucleotide analogs is highly dependent on their chemical state. Future studies should focus on identifying the specific degradation products responsible for the observed toxicity and elucidating the underlying mechanisms of action.

**Figure 6.**
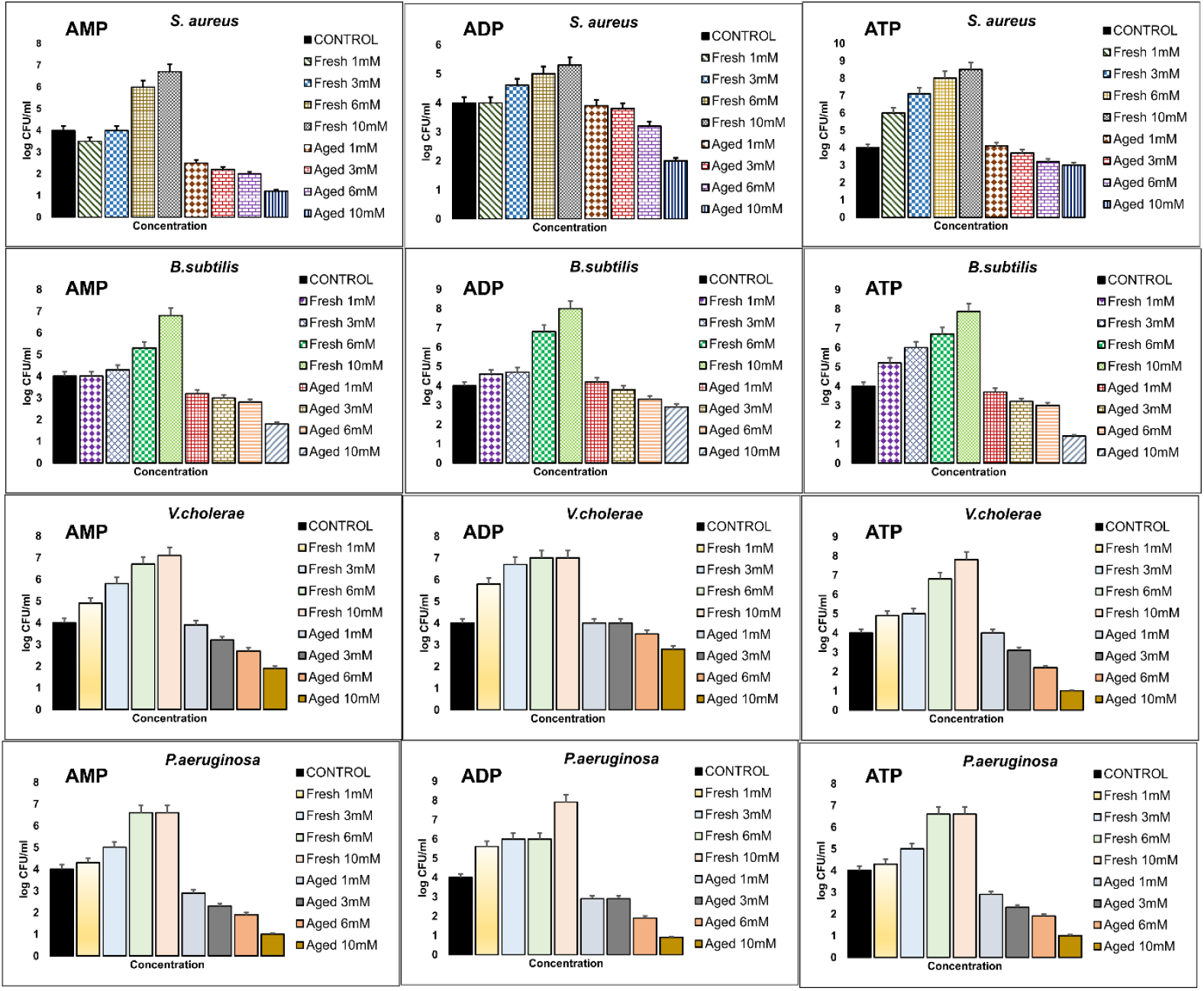
Antibacterial study graph of Nucleotides on four gram-positive and gram-negative strains.

## Conclusions

The self-assembling properties of AMP, ADP, and ATP have been studied in detail by different microscopy methods. The self-assembly studies suggest aggregation of AMP, ADP, and ATP is concentration as well as time dependent. The SEM and AFM studies suggest small globular structures fuse with each other after ageing and may form small fibrils. Thioflavin T assay suggests the amyloid nature of the assemblies and the toxicity of aggregates. Hence, the results presented may have implications in understanding the patho-physiology of cancer, neurodegenerative diseases inflammation, and many other diseases associated with the accumulation of these nucleotides from an amyloid perspective and possibly pave the way for common therapeutic strategies for treating these diseases. In the future, we aim to conduct additional research on these nucleotides, drawing insights from our group’s current and previous studies. We wish to perform extensive biophysical and crystal growth assays to characterize the molecular nature of the amyloid formation from these nucleotides.

## Experimental Section

Research on self-assembly: AMP, ADP and ATP were purchased from Sigma-Aldrich without further purification (99% pure). JSK Fine chemicals provided solvents, while Millipore instrument supplied deionized water. All solvents utilized were of analytical grade and had a purity of 99%. These solutions were left to incubate at various time intervals at room temperature. Each sample underwent optical microscopy analysis by dropping 20μL onto a glass slide and letting it dry. The Leica DM2500 microscope was used to capture images at 40X and 63X magnification once the samples were dried. The SEM images were taken using a Nova Nano FEG-SEM 450 microscope at an accelerating voltage range of 5 to 15 kV. Silicon wafers were employed in the preparation of the SEM samples. FEI Technai 20 U Twin Transmission Electron Microscope was utilized for TEM analyses, with the samples being dried on copper grids coated with carbon on a 400 mesh. Asylum Research OXFORD atomic force microscopy (MFP3D) with ARC2 controller was utilized for AFM analysis. The research utilized AFM tips (AC240TS-R3) sourced from Asylum Research Probes, featuring a radius close to 7 nm. AFM samples were made by dropping and drying 10μL of aqueous nucleotides solution. In the coincubation experiment, the nucleotides were co-incubated with urea and tannic acid in a 1:1 ratio. TA and urea with a 99% purity were acquired from Sigma-Aldrich and Avra chemicals ThT assay of nucleotides in solution were done using a Cary Eclipse Fluorescence Spectrophotometer, the fluorescence spectra were captured. With a 5 nm excitation and emission slit, all of the data were captured at an excitation wavelength of 450 nm. ThT used in the studies was of 99 percent purity purchased from SRL. The sample preparation of microscopy and spectroscopy were done by previously reported methodologies used in our Lab.

### Cell Culture

Human retinal pigment epithelial cells (RPE-1) and human colorectal carcinoma cells (HCT116) were obtained from a certified cell bank. Both cell lines were cultured in Dulbecco’s Modified Eagle’s Medium (DMEM) supplemented with 10% fetal bovine serum (FBS), 100 U/mL penicillin, and 100 μg/mL streptomycin at 37°C in a humidified atmosphere of 5% CO_2_. Cells were subcultured every 3-4 days using trypsin-EDTA solution when reaching 70-80% confluence. Adenosine monophosphate (AMP), adenosine diphosphate (ADP), and adenosine triphosphate (ATP) were purchased from a commercial supplier. Stock solutions of AMP, ADP, and ATP were prepared in phosphate-buffered saline (PBS) at concentrations of 100 mM, then filtered through a 0.22 μm membrane to ensure sterility. These stock solutions were diluted to working concentrations of 1 mM, 3 mM, 6 mM, and 10 mM in cell culture media immediately prior to use. For the aged conditions, AMP, ADP, and ATP solutions were incubated at 37°C for 72 hours to induce aggregation and simulate the aged state, verified by spectroscopic analysis. Freshly prepared and aged solutions were used in parallel for all experiments. Cell viability was assessed using the MTT assay to evaluate the effect of AMP, ADP, and ATP in both fresh and aged conditions on RPE-1 and HCT116 cells. The MTT assay was performed in 96-well plates with the following experimental groups: RPE-1 cells were seeded in 96-well plates at a density of 10,000 cells per well. After 24 hours, cells were treated with fresh or aged AMP at concentrations of 1 mM, 3 mM, 6 mM, and 10 mM. Control wells were treated with PBS alone. Cells were incubated for 24 hours, after which cell viability was measured by adding 10 μL of 5 mg/mL MTT solution to each well. The plates were incubated at 37°C for 4 hours. The resulting formazan crystals were solubilized with 100 μL of DMSO, and absorbance was measured at 570 nm using a microplate reader. HCT116 cells were seeded under the same conditions as RPE-1 cells and treated similarly with fresh and aged AMP. The MTT assay was performed as described above to evaluate cell viability. RPE-1 cells were seeded and treated with fresh or aged ADP at 1 mM, 3 mM, 6 mM, and 10 mM concentrations. After a 24-hour incubation, the MTT assay was conducted as previously described to determine the effect of ADP on cell viability in both conditions. HCT116 cells were treated with ADP in a similar manner as RPE-1 cells, and the MTT assay was performed to assess the cytotoxicity or proliferative effect of fresh and aged ADP on this cell line. RPE-1 cells were exposed to fresh and aged ATP at concentrations of 1 mM, 3 mM, 6 mM, and 10 mM for 24 hours. Cell viability was determined using the MTT assay, as described above. HCT116 cells were similarly treated with fresh and aged ATP, and their viability was measured using the MTT assay to compare the effects of ATP in both conditions. The absorbance data from the MTT assay were normalized to untreated control cells, and the results were expressed as a percentage of viable cells. Each experimental condition was performed in triplicate, and all experiments were repeated three times. Statistical analysis was carried out using one-way ANOVA followed by Tukey’s post-hoc test for multiple comparisons. Data were presented as mean ± standard deviation (SD), and a p-value of <0.05 was considered statistically significant.

### Antimicrobial Assay via Broth Microdilution Method

Four different strains of bacteria were selected for this investigation: two Gram-positive species (*Bacillus subtilis* and *Staphylococcus aureus*) and two Gram-negative species (*Pseudomonas aeruginosa* and *Vibrio cholerae*). The strains were initially grown in Mueller-Hinton (MH) broth and then incubated at 37°C with constant shaking (150 rpm) until the strains reached an optical density (OD) of 0.4 at 600 nm (OD600), which is indicative of the mid-logarithmic phase of bacterial growth. Using serial dilution and plating, the bacterial density at an OD600 of 0.4 was found to be approximately 1 × 10^8^ CFU/mL. After bacterial cultures were established at OD600 = 0.4, 10 mL aliquots of each culture were transferred into sterile tubes. A final volume of 10 mL per sample was maintained by adding test samples (ATP, ADP, and AMP) in equal concentrations of 1mL to the aliquoted bacterial suspensions that had been prepared at different concentrations (1mM, 3mM, 6mM, and 10mM). In order to evaluate baseline growth and ensure the assay conditions were sterile, controls such as media-only controls and untreated bacterial cultures were prepared concurrently. All bacterial cultures, including treated and control samples, were incubated at 37°C with shaking conditions for a period of 24 hours. After incubation, each bacterial culture’s optical density (OD600) was measured using a spectrophotometer. Bacterial growth was assessed by comparing the OD600 values of cultures treated with different concentrations (1mM, 3mM, 6mM, and 10mM) of the test samples to those of untreated control cultures. The reduction in OD600 indicated the inhibitory effect of the test samples on bacterial growth in a concentration-dependent manner. The optical density (OD600) of the treated bacterial cultures was examined for each concentration of the test samples after a 24-hour incubation period. To investigate if varying concentrations affected bacterial growth, the OD600 values were plotted directly against the appropriate concentrations of the test compounds. The concentration vs OD600 plot gave a concise representation of the antimicrobial activity, where lower OD values indicated higher levels of bacterial growth inhibition [21].

## Acknowledgments

Dr. Nidhi Gour (NG), RD greatly acknowledge support from SERB research grant SERB SPG/2021/000521 for funding and fellowships and Indrashil University for infrastructure support. KP and RP acknowledge support from Indrashil University for infrastructure support. AS thanks DST for PMRF fellowship, DB thanks SERB-CRG, MoES-STARS, Gujcost-DST, GSBTM and IITGN for research grants.

