## Supporting information for "Pathophysiological Implications of Nucleotide Self-Assembly: Adenine-Derived Nucleotides Aggregation in Disease Mechanisms"

| **Sr.No.** | **Topic** | **Page No.** |
| --- | --- | --- |
| 1. | Experimental procedures  Optical microscopic studies | 2 |
| 2. | Cross seeding Experiments | 2-3 |

**Experimental procedures:**

**Optical microscopic studies:**

The Nucleotides were purchased from Sigma- Aldrich without further purification (99% pure). The sample preparations were done in deionized water. These solutions were incubated for varied amount of time at 37.4 C. Each sample was spread out over a glass slide in 20 µL portions for OM imaging, and then it dried. The images were taken using Leica DM2500 microscope under 40X and 63X magnifications.

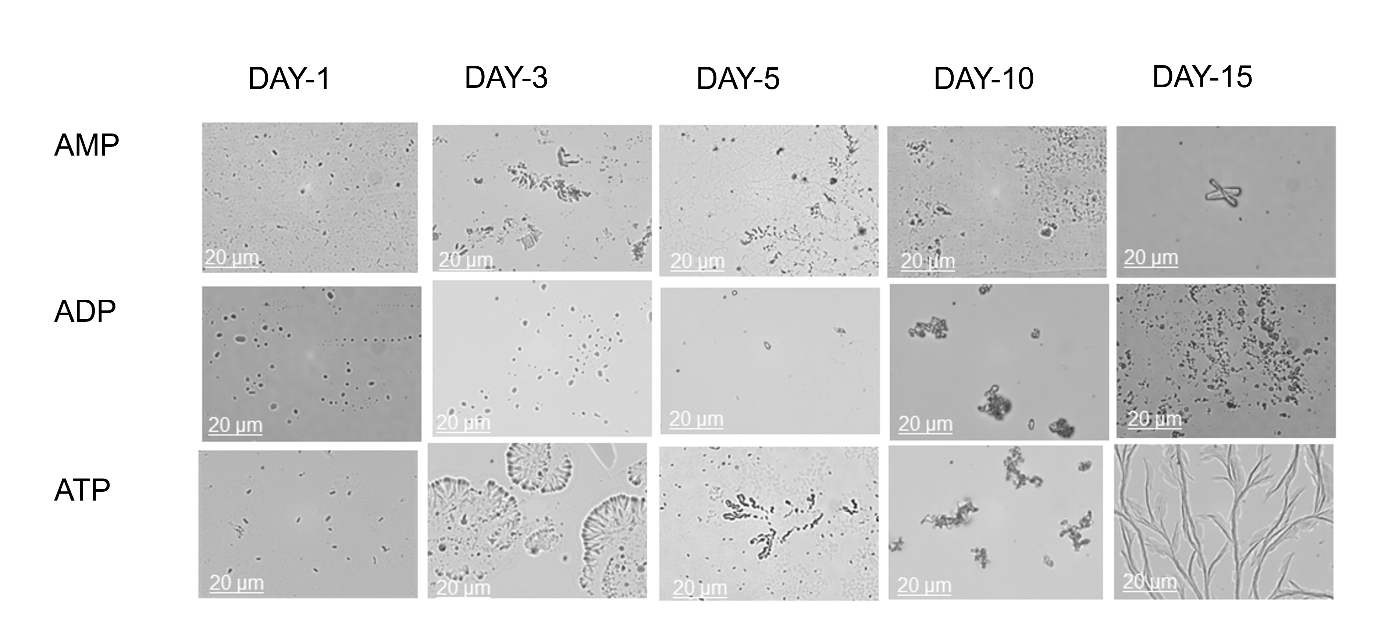

**Figure S-1:** AMP, ADP and ATP at various time intervals cross-seeded with urea in 1:1 proportion.

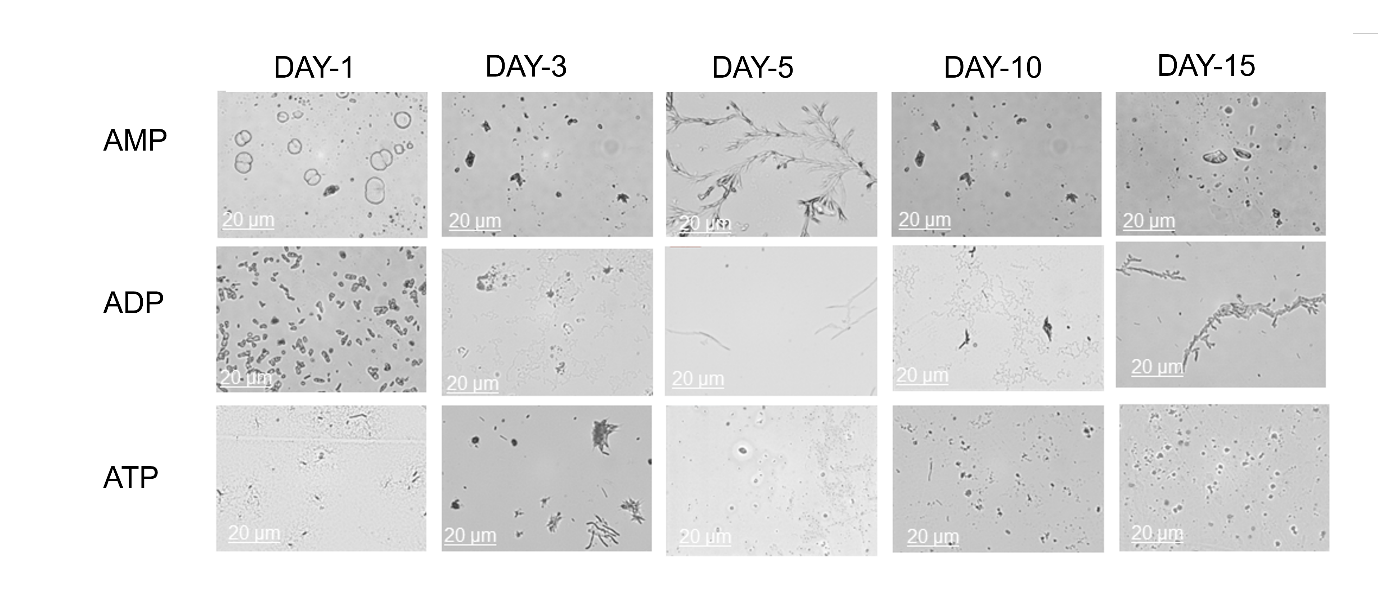

**Figure S-2:** AMP, ADP and ATP at various time intervals cross-seeded with Tannic acid in 1:1 proportion.

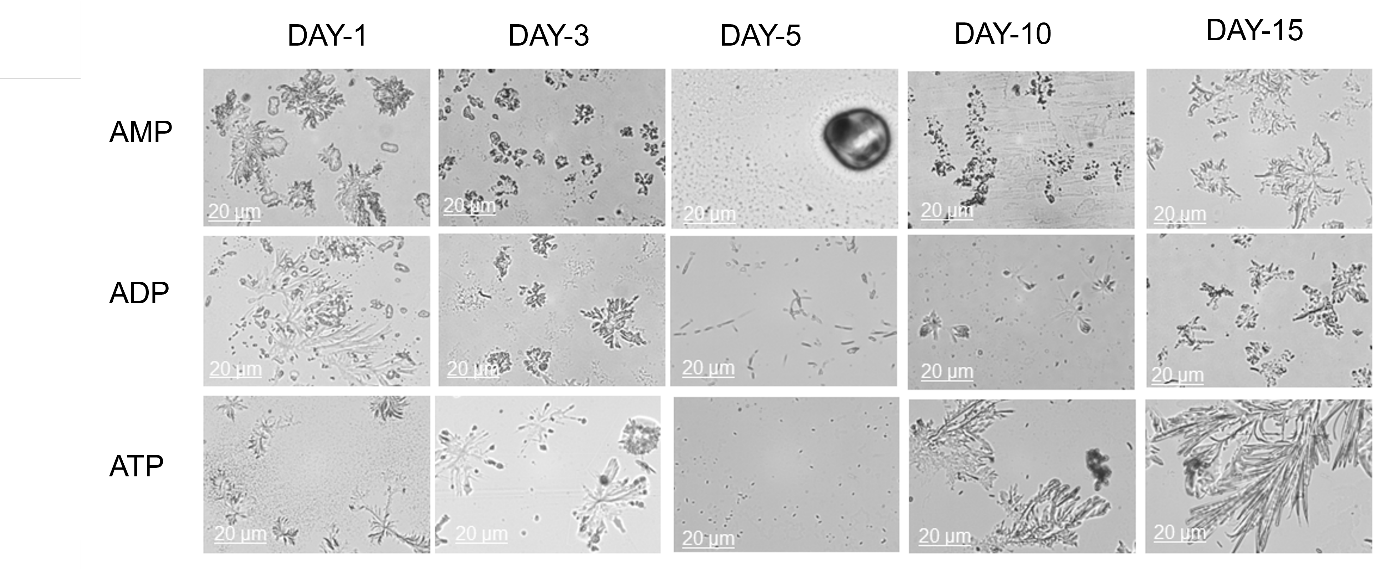

**Figure S-3:** AMP, ADP and ATP at various time intervals cross-seeded with Phenylaniline in 1:1 proportion.
